# AutoBlot: deterministic single-cell western blotting reveals proteomic diversity in rare cell populations

**DOI:** 10.64898/2026.06.22.733890

**Authors:** Cyril Deroy, Md Nazibul Islam, Nadine Goldhammer, Lorraine Jiang, Jennifer M. Rosenbluth, Amy E. Herr

## Abstract

Protein abundance and proteoform composition are direct determinants of cellular phenotype, yet both remain difficult to quantify in individual cells from specimens available in limited numbers. Single-cell western blotting (scWB) provides quantitative, molecular-mass-resolved protein measurements but relies on Poisson-limited gravity settling, requiring approximately 10⁶ starting cells while constraining single-cell microwell occupancy to a theoretical maximum of 36.8%. Here, we introduce AutoBlot, an image-guided piezoelectric dispensing approach that deterministically loads individual cells into predefined scWB microwells. AutoBlot achieved approximately 98% single-cell occupancy among targeted microwells while operating from as few as 10,000 starting cells. Across the tested cell concentrations, measured occupancy exceeded Poisson predictions by 31- to 904-fold, and the same dispensing strategy enabled deterministic cell-bead co-loading. We applied AutoBlot to 1,449 cells from seven patient-derived breast organoids generated from histologically normal mammary tissue. Measurements of five lineage-associated proteins revealed donor-to-donor variation, differences associated with germline BRCA1 and menopausal status, and discordance between surface-marker- and cytokeratin-based lineage assignments. AutoBlot also resolved ER-immunoreactive species at approximately 66 and 46 kDa in individual PDO cells. These results establish deterministic cell loading as a strategy for extending molecular-mass-resolved single-cell protein analysis to heterogeneous, cell-limited specimens.

## Introduction

While single-cell sequencing tools have matured rapidly, targeted single-cell protein assays remain limited yet essential for hypothesis-driven research and biomarker studies^1^. As a targeted proteomic tool, the single-cell western blot (scWB) assay simultaneously quantifies protein expression across hundreds of individual cells with molecular-mass specificity^2,3^. By integrating polyacrylamide gel electrophoresis (PAGE) for size-based protein separation with in-gel immunoprobing, scWB confers selectivity advantages over antibody-binding-only methods, including the ability to resolve protein isoforms and distinguish immunoreactive species according to apparent molecular mass^2^. This selectivity is not available to antibody-binding-only platforms such as flow or mass cytometry, in which all immunoreactive species collapse into a single per-cell intensity value. Since its introduction, the platform has been extended to circulating tumor cells^4^, protein complexes^5^, three-dimensional projection electrophoresis^6^, multimodal nucleic acid and protein isoform profiling^7^, and chemically fixed biorepository specimens^8^. Across these implementations the separation chemistry, immobilization strategy, and immunoprobing workflow have advanced substantially, whereas cell isolation to the microwell array has remained largely unchanged. Although microtransfer pipette-based seating has demonstrated deterministic single-cell loading at 100% efficiency from ultra-low inputs, including individual blastocysts and blastomeres4,7,9, this approach requires exceptional manual dexterity, is inherently low-throughput, and does not scale to the hundreds of cells per chip needed for population-level proteomic profiling. The majority of scWB implementations instead rely on Poisson-distributed gravity settling to load cells into microwells, resulting in single-cell occupancy in only 10–30% of microwells and requiring ≥10⁶ starting cells to populate enough microwells for statistically powered analyses. The ceiling here is statistical rather than technical: for stochastic loading of a microwell of volume V from a suspension of concentration C, the probability of exactly one cell per well follows P(X = 1) = λe⁻λ with λ = CV, which is maximized at 36.8% when λ = 1. Raising the input concentration to fill more wells necessarily raises the multi-cell occupancy that single-cell resolution forbids. This constraint restricts scWB to abundant cell populations and excludes rare or precious specimens, such as patient-derived organoids (PDOs), fine needle aspirates, and circulating tumor cells, where cell numbers are limited to 10³–10⁵. PDOs exemplify this specimen class: they are inherently cell-limited (typically 10³ to 10⁵ cells per 50 µL dome), encompass multiple epithelial lineages, and enable benchmarking against well-characterized mammary biology^14^.

Here we report AutoBlot, which replaces Poisson-limited gravity settling with piezoelectric dispensing under real-time imaging feedback to achieve deterministic loading of scWB microwell arrays. AutoBlot attains approximately 98% single-cell occupancy from a starting suspension of 10,000 cells, a 100-fold reduction in starting cell requirement relative to gravity settling, and extends the same deterministic control to co-loading of one cell and one bead per microwell. We applied AutoBlot to 1,449 single cells from seven patient-derived breast organoids established from histologically normal mammary tissue, quantifying five lineage-associated proteins and resolving ER-immunoreactive species at approximately 66 and 46 kDa. The measurements revealed donor-to-donor variation, differences associated with germline BRCA1 status and menopausal status, and discordance between surface-marker- and cytokeratin-based lineage assignments.

## Results and discussion

### Deterministic single-cell and cell–bead loading of scWB microwell arrays

We developed AutoBlot, which replaces gravity settling with the CellenONE robotic dispenser for deterministic cell loading into scWB microwell arrays (Fig. 1a,b). Previously used to dispense single cells into 384-well plates and nanoliter droplets^10,11^, we adapted the CellenONE to deposit individual cells into sub-100-µm microwells using real-time imaging and piezoelectric actuation. Starting from a suspension containing 10,000 cells, AutoBlot deterministically loaded 1,000 single cells into preselected microwells with approximately 98% targeted-well occupancy (Fig. 1a, SI Fig. 1). Here, targeted-well occupancy denotes the fraction of programmed microwells containing exactly one cell. Across input concentrations of 10,000, 30,000, and 300,000 cells mL−1, the measured single-cell occupancy exceeded the corresponding Poisson predictions by 904-, 302-, and 31-fold, respectively (Fig. 1c). Moreover, deterministic co-loading of one cell and one bead at concentrations of 30,000 mL⁻¹ each, achieved comparable targeted-well occupancy and exceeded the Poisson-predicted probability of one-cell–one-bead co-occupancy by approximately five orders of magnitude (Fig. 1c). Together, these measurements demonstrate that coordinate-defined deposition circumvents the Poisson trade-off between singlet occupancy and multiplet formation while reducing the starting-cell requirement from approximately 10⁶ to 10⁴ cells. Because AutoBlot modifies the cell-loading step while retaining downstream PAGE separation, protein immobilization, and immunoprobing, we applied the workflow to patient-derived breast organoids (PDOs) to evaluate molecular-mass-resolved single-cell protein analysis in a specimen comprising limited cell numbers and multiple mammary epithelial lineages. Lysis buffer composition and electrophoresis duration were optimized for PDO solubilization (SI Fig. 2), and Table 1 summarizes the range of formulations employed across scWB publications^2–8,12,13^ providing a consolidated reference for the field.

**Fig. 1:**
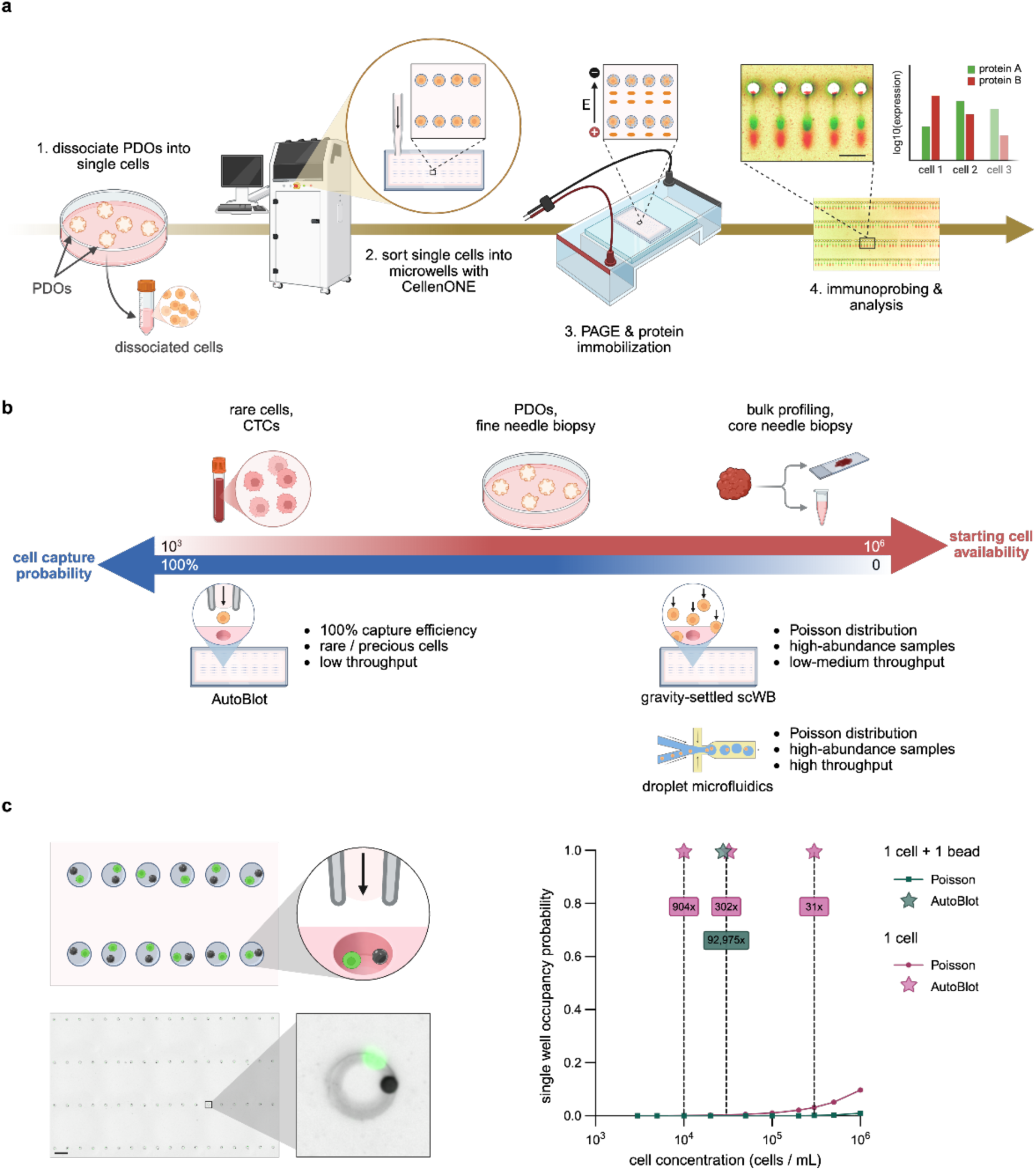
AutoBlot achieves deterministic scWB of cell-limited specimens. **a**, Schematic of the AutoBlot workflow. PDOs are enzymatically dissociated into single-cell suspensions and loaded into the CellenONE robotic dispensing system, which deterministically deposits individual cells into polyacrylamide microwells (48 µm diameter × 60 µm height) with approximately 98% single-cell occupancy among addressed microwells while operating from a starting suspension of 10,000 cells. Polyacrylamide gel electrophoresis separates proteins by molecular mass, UV-initiated immobilization fixes separated proteins in the gel, and immunoprobing with fluorescently labeled antibody probes detects a panel of proteins across thousands of single cells simultaneously. A representative bar graph (right) illustrates per-cell quantification of the 5 protein panel from the scWB array (scale bar = 300 µm). **b**, Comparison of single-cell isolation strategies. A spectrum of platforms is shown ranging from high cell-capture probability with low starting cell requirements (left) to low capture probability requiring large cell inputs (right). AutoBlot (CellenONE-based) achieves approximately 98% single-cell occupancy from 10^4^ starting cells, thus corroborating suitability for rare or precious cells. At intermediate starting cell availability (10⁴), AutoBlot is well suited for PDOs and fine needle biopsies. Gravity-settled scWB and droplet microfluidics offer higher total cell throughput but depend on Poisson-distributed loading and require 10⁶ starting cells, making them appropriate for bulk profiling. **c**, Deterministic co-encapsulation of single cells and beads into microwells. Left: schematic (top) illustrating CellenONE-based sequential dispensing of a single cell (green) and a single bead (black) into each polyacrylamide microwell of the scWB chip. Fluorescence and brightfield micrograph (bottom) of the scWB microwell array after co-loading, showing a bead-cell pair in each well (scale bar = 300 µm); inset shows a magnified view of a single microwell containing one cell and one bead. Right: graph comparing AutoBlot co-encapsulation efficiency against the theoretical Poisson distribution. Lines show the Poisson probability of capturing exactly one cell per well (pink) or one cell and one bead per well (teal) as a function of input cell concentration. Stars mark the cell concentrations tested experimentally with AutoBlot, at which single-cell loading exceeded Poisson probability by 904×, 302×, and 31×, and co-encapsulation loading exceeded Poisson probability by 92,975×.

**Fig. 2.**
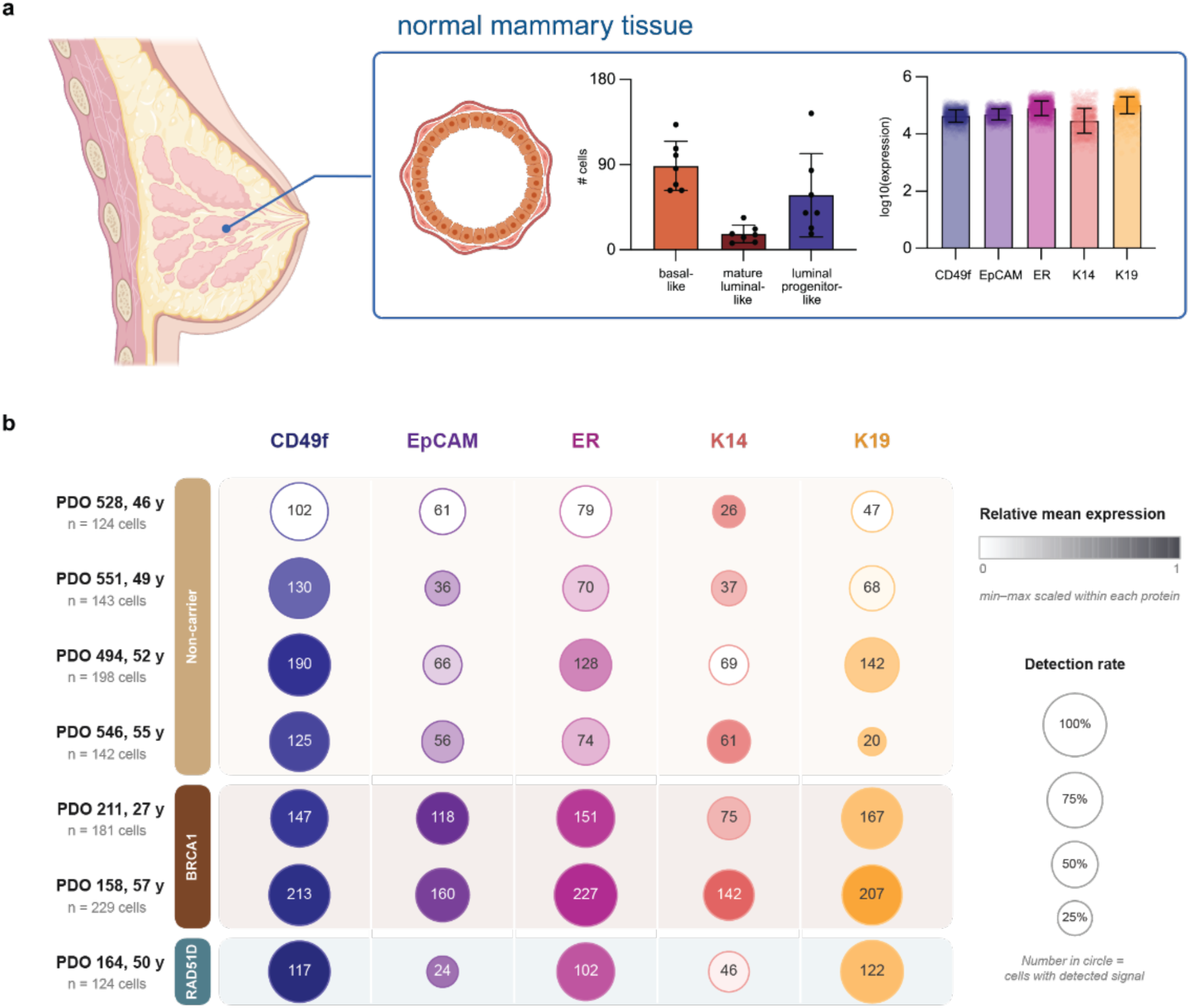
| AutoBlot resolves basal and luminal marker expression in single cells and reveals donor-to-donor variation across normal mammary organoids. **a**, Protein profiling of normal mammary tissue. Non-carrying mammary tissue displays organized ductal architecture with distinct basal myoepithelial and luminal epithelial layers. Five panel proteins were measured across 7 normal mammary tissue PDOs: basal markers CD49f and K14, luminal markers EpCAM and K19, and estrogen receptor (ER). Left bar graph: mean cell-type composition defined by surface markers CD49f and EpCAM (points represent per-PDO cell numbers). Right bar graph: mean protein expression levels (points represent individual cells). Error bars represent standard deviation. **b**, Single-cell protein detection and expression across the PDO cohort. Bubble matrix summarizing scWB results for the same five protein targets across all 7 normal mammary tissue PDOs (n = 1,141 cells with an assigned cell type; a further 308 assayed cells could not be lineage-classified and are excluded here, so detection rates refer to lineage-classified cells). Each circle represents one organoid–protein combination. Circle area is proportional to the detection rate (the fraction of cells in which the protein was detected above the limit of detection); the number inside each circle gives the absolute count of cells with detected signal, and the total number of cells assayed per organoid is given in the row labels (n). Circle shading indicates the mean log10(Expression + 1) among detected cells, normalized independently within each protein column (per-protein min–max scaling across organoids; 0 = lowest mean expression, 1 = highest); each column is shaded in the color used for that marker in a. Organoids are grouped by germline status: non-carrier (tan), BRCA1 (dark brown) and RAD51D (teal): ordered by donor age within each group, and labeled with donor age. Detection rate and expression data were derived from automated fluorescence image analysis of scWB arrays as described in Methods.

**Table 1.** Lysis buffer formulations for scWB have evolved from standard RIPA to urea-supplemented compositions for challenging sample types. Concentrations are reported as weight/volume (wt/v) or volume/volume (v/v) fractions. Buffer components include SDS (sodium dodecyl sulfate), Triton X-100 (non-ionic surfactant), Na-DOC (sodium deoxycholate), and urea (chaotropic agent). The rightmost columns reflect recent formulations that incorporate 8 M urea for enhanced protein solubilization of PDO-derived and fixed cells. The AutoBlot formulation uses 2× RIPA + 8 M urea (1% SDS, 0.5% Na-DOC, 0.2% Triton X-100, 2× Tris-glycine).

| Component | Std. RIPA Buffer | Hughes et al., 2014 | Kang et al., 2016 | Duncombe et al., 2016 | Sinkala et al., 2017 |  |  | Zhang et al., 2019 | Grist et al., 2020 |  |  | Vlassakis et al., 2021 | Kim et al., 2021 | Liu and Herr, 2023 |  |  |
| --- | --- | --- | --- | --- | --- | --- | --- | --- | --- | --- | --- | --- | --- | --- | --- | --- |
|  |  |  |  |  | P1 (incomplete solubilization) | P2, 3 & 4 | P5, 6 & 7 |  | 1× RIPA 1× Tris-gly | 2× RIPA 2× Tris-gly | Modified 2× RIPA 2× Tris-gly |  |  | AR buffer (live cells) | AR buffer (fixed cells) | Running buffer |
| SDS (wt/v) | 0.10% | 0.50% | 0.50% | 1% | 0.50% | 1% | >1% | 1% | 0.50% | 1% | 1% | 0.50% | 1% | 0.50% | 1% | 1% |
| Triton X-100 (v/v) | — | 0.10% | 0.10% | 0.10% | 0.10% | 0.25% | 1% | 0.10% | 0.10% | 0.20% | 0.20% | 0.10% | 0.10% | — | — | — |
| Na-DOC (wt/v) | 1% | 0.25% | 0.25% | 1% | 0.25% | 0.25% | 0.50% | 0.50% | 0.25% | 0.50% | 0.50% | 0.25% | — | — | — | — |
| Tris | — | 12.5 mM | — | — | — | — | — | — | — | — | — | — | — | — | — | — |
| Glycine | — | 96 mM | — | — | — | — | — | — | 0.5× | — | — | — | — | — | — | — |
| Tris-glycine | — | — | 0.5× | 1× | 1× | 1× | 1× | — | 1× | 2× | 2× | 0.5× | 1× | 1× | 1× | 1× |
| Tris-HCl | 25 mM | — | — | — | — | — | — | — | — | — | — | — | — | — | — | — |
| NaCl | 150 mM | — | — | — | — | — | — | — | — | — | — | — | — | — | — | — |
| NaF (phosphoprotein) | — | 1 mM | — | — | — | — | — | — | — | — | — | — | — | — | — | — |
| Na<sub>3</sub>VO<sub>4</sub> (phosphoprotein) | — | 1 mM | — | — | — | — | — | — | — | — | — | — | — | — | — | — |
| NP-40 | 1% | — | — | — | — | — | — | — | — | — | — | — | — | — | — | — |
| Urea | — | — | — | — | — | — | — | — | — | — | 8 M | 8 M | 8 M | 6 M | 6 M | — |
| pH | 7.6 | 8.3 | — | — | — | — | — | — | — | — | — | — | — | — | — | — |

Abbreviations: SDS, sodium dodecyl sulfate; Na-DOC, sodium deoxycholate; NP-40, Nonidet P-40; AR, antigen retrieval; Std., standard; Tris-gly, Tris-glycine. Concentrations for SDS, Triton X-100, Na-DOC, and NP-40 are expressed as percentages. Em dashes (—) indicate that the component was not included in the buffer formulation.

### Targeted single-cell protein profiling reveals donor-to-donor variation across normal breast PDOs

We applied AutoBlot to profile the targeted single-cell proteomic landscape of patient-derived breast organoids from 7 donors with histologically normal mammary tissue (Fig. 2a). A panel of five proteins was measured across 1,449 single cells from the 7 donors: CD49f (∼125 kDa; UniProt: P23229), K14 (∼50 kDa; UniProt: P02533), EpCAM (∼40 kDa; UniProt: P16422), K19 (∼44 kDa; UniProt: P08727) and estrogen receptor (ER) (∼66 kDa full-length ERα; UniProt: P03372). CD49f and EpCAM are established surface markers used to distinguish basal/myoepithelial, mature luminal, and luminal progenitor mammary epithelial populations^14^, whereas K14 and K19 are intracellular cytokeratins associated with basal and luminal identity, respectively^15^. ER identifies a hormone-responsive luminal subset^13^. Together, these markers enabled parallel lineage assignments based on CD49f/EpCAM and K14/K19 measurements. Proteins were assayed after PAGE separation, enabling multiplexed surface and intracellular protein measurements from the same cells. For each protein marker, both the fraction of positive cells and the mean expression among positive cells varied across PDOs (Fig. 2b). In a qualitative reference comparison, an independent breast cancer PDO exhibited a broader cell-to-cell distribution of EpCAM signal intensity than the MCF-7 cell line (SI Fig. 3), consistent with the greater phenotypic heterogeneity expected in a primary organoid culture. Of the 1,449 assayed cells, 1,141 cells with an assigned lineage were included in the cohort-level detection matrix, whereas 308 cells that could not be assigned a lineage were excluded from this visualization. For each PDO–protein combination, the matrix separately reports the fraction of cells with signal above the detection threshold and the mean expression among detected cells. This analysis therefore distinguishes variation in the frequency of protein detection from variation in signal magnitude among protein-positive cells.

**Fig. 3.**
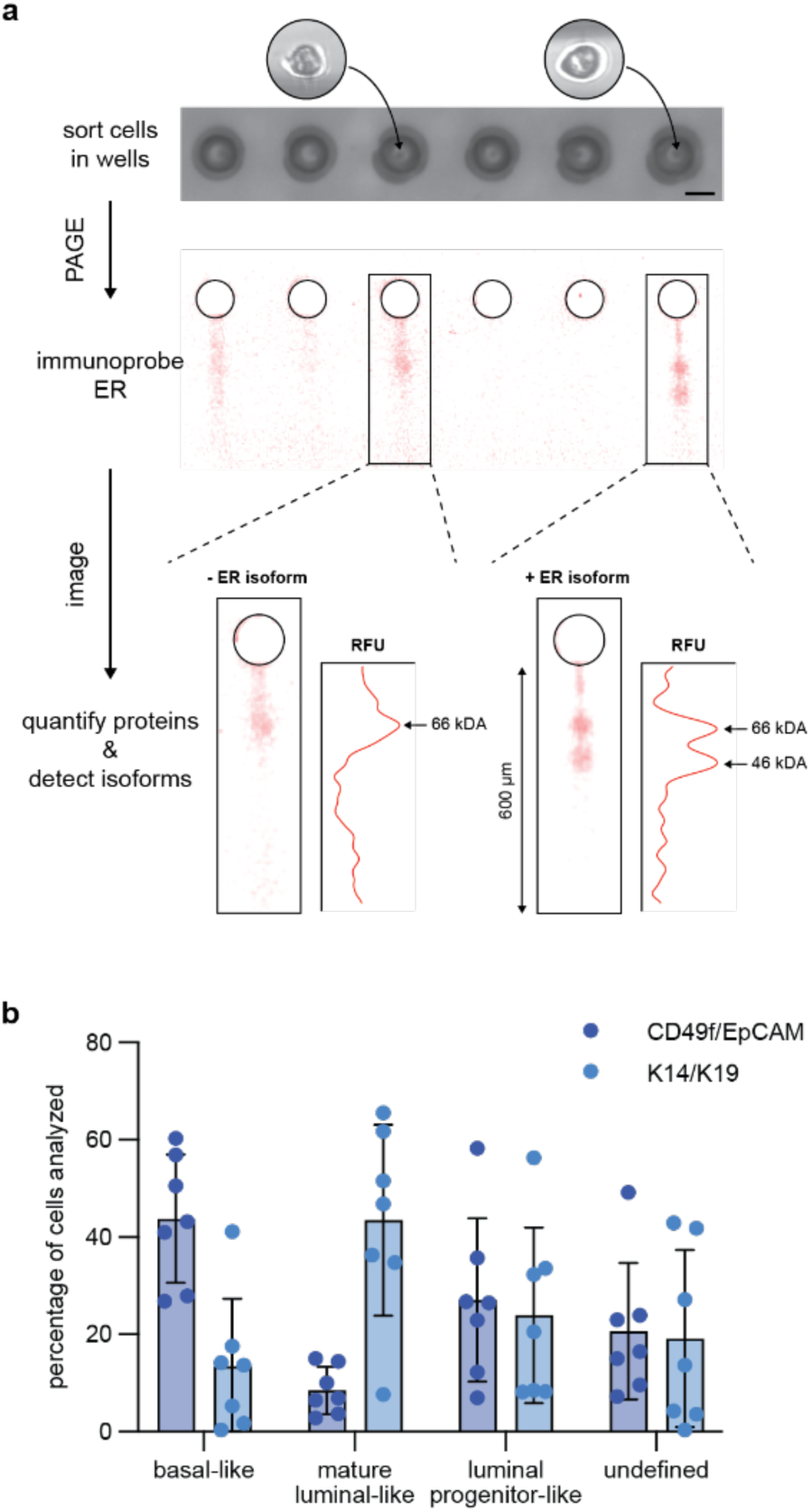
| AutoBlot resolves ER isoforms in single cells and reveals discordance between surface and cytokeratin lineage markers. **a**, scWB array and ER isoform detection. Top: brightfield micrograph of the scWB microwell array (6 microwells shown) after sorting cells with the CellenONE (scale bar = 100 µm). Sorted cells are visible as faint black spots in the center of each microwell. (Inset) Brightfield micrographs collected during CellenONE cell dispensing show sorted cells (∼25 µm in diameter) in wells 3 and 6. Middle: fluorescence micrograph of the scWB microwell array shows scWB-resolved ER protein bands separated by PAGE. Bottom: magnified views of representative single-cell PAGE lanes. Left lane: a single ER protein band at ∼66 kDa (full-length ERα). Right lane: two resolved proteins at ∼66 kDa and ∼46 kDa, demonstrating resolution of ER isoforms at single-cell resolution. The ∼46-kDa isoform corresponds to a truncated ERα variant lacking part of the activation function-1 domain. Fluorescence intensity profiles (relative fluorescence units) accompany each scWB lane, with arrows indicating molecular mass positions. **b**, Discordance between surface and intracellular lineage markers. Bar graphs comparing cell-type distributions from CD49f/EpCAM surface marker classification (navy) versus K14/K19 cytokeratin classification (light blue) across normal mammary tissue PDOs (n = 7), with surface-marker classification assigning a greater proportion of cells to the basal-like category than cytokeratin classification in every PDO. The “undefined” category comprises cells in which only ER was detected and the remaining markers were below the detection threshold, precluding lineage classification. Points represent individual PDOs.

### AutoBlot resolves ER-immunoreactive species and discordant lineage-marker assignments

A key advantage of scWB over antibody-binding-only immunoassays is the ability to resolve proteins by molecular mass. Leveraging this capability, we observed two distinct ER-immunoreactive bands at approximately 66 and 46 kDa in a small subset of individual PDO cells (Fig. 3a), despite the stringent lysis conditions required for PDO solubilization. The apparent molecular masses of these bands are consistent with full-length ERα66 and the truncated ERα46 isoform reported in previous bulk and single-cell immunoblotting studies^13,16^. ERα46 lacks the N-terminal activation function-1 domain and exhibits functional properties distinct from ERα66; reduced ERα46 expression has also been associated with secondary tamoxifen resistance in a subset of breast cancer patients^16^. The anti-ERα antibody used in this experiment (clone SP1) is directed against a C-terminal epitope common to both ERα66 and ERα46, consistent with detection of both species by the same probe within a single immunoprobing round. Previous scWB studies resolved ERα66 and ERα46 and characterized heterogeneous responses of these isoforms to tamoxifen in breast cancer cell lines^13^; AutoBlot extends molecular-mass-resolved detection of these ER-immunoreactive species to cell-limited PDO specimens. Because the ∼46-kDa species was observed in only a small subset of cells (>2%) within a given PDO, its signal in a bulk lysate would be proportionally diluted relative to the dominant ∼66-kDa species and could fall below bulk detection, highlighting the value of resolving molecular-mass heterogeneity at single-cell resolution.

Beyond resolving molecular-mass heterogeneity, AutoBlot also enables multiple protein markers to be measured within the same cell, allowing direct comparison of alternative phenotypic classification schemes. Although CD49f/EpCAM and cytokeratin staining are widely used to characterize mammary lineages^17,18^, it remains unclear to what extent lineage assignments derived from these marker sets agree at the single-cell level. The ability of scWB to simultaneously measure surface and intracellular proteins from the same cell enabled this comparison at the single-cell protein expression level across 7 normal breast PDOs. Cells could be assigned into basal-like, mature luminal-like, and luminal progenitor-like categories using either classification scheme. Surface-marker-based classification consistently identified a greater proportion of basal-like cells than cytokeratin-based classification across all PDOs (Fig. 3b). The systematic difference between the two classification schemes is consistent with their capturing distinct aspects of mammary epithelial state rather than interchangeable lineage identities. CD49f/EpCAM report cell-surface phenotype, whereas cytokeratins reflect intracellular epithelial differentiation, and both marker classes can exhibit context-dependent expression^19^. This distinction may be particularly relevant in the organoid setting, where CD49f (integrin α6) engages laminin^20^, a major component of the basement membrane extract used for PDO propagation^21^.

### Single-cell protein phenotypes vary with germline BRCA1 status, menopausal status, and donor identity

Among the 7 normal-tissue PDOs, we excluded one RAD51D mutation carrier and compared the remaining 6 donors: 2 BRCA1 germline mutation carriers versus 4 non-carrier controls (Fig. 4a). BRCA1-carrier PDOs showed a markedly expanded luminal progenitor population (55.6% versus 26.2% in non-carrier samples), consistent with previous studies reporting increased luminal progenitor representation in mammary tissue and organoid cohorts from BRCA1 mutation carriers^15,22–24^. This concordance with established BRCA1-associated mammary biology supports the biological interpretability of the marker-defined populations measured by AutoBlot. The luminal progenitor compartment is of specific interest in this genotype because it has been proposed as the candidate cell of origin for BRCA1-associated basal-like tumours^22,23^, making its representation in histologically normal, cancer-prone tissue a relevant quantity to measure directly. AutoBlot further quantified per-cell protein expression levels, with higher mean EpCAM and ER signals observed in the two BRCA1-carrier PDOs than in the four non-carrier PDOs (Fig. 4a). CD49f, K14, and K19 showed smaller between-group differences in this cohort. Because both PDO groups were classified using the same CD49f/EpCAM scheme, the comparison in Fig. 4a is internally controlled with respect to the scheme-dependent differences described above.

**Fig. 4.**
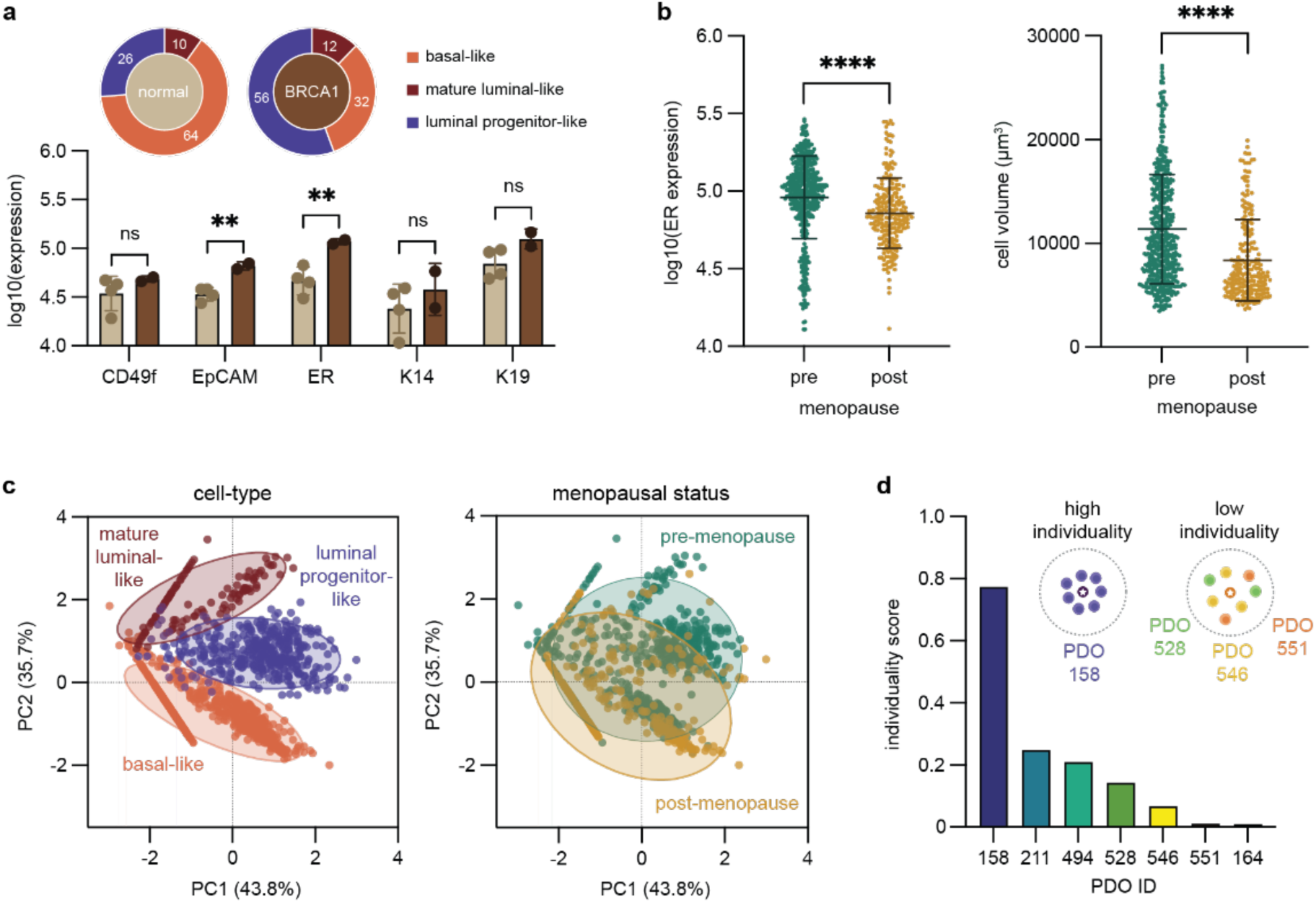
| Single-cell protein phenotypes are associated with germline BRCA1 status and menopausal status in mammary organoids. **a**, BRCA1 genotype-associated differences. Top: donut charts comparing cell-type composition (percentage of cells analyzed) between non-carriers (n = 4 donors) and BRCA1-carrier (n = 2 donors) PDOs, colored by cell type (basal-like, coral; mature luminal-like, dark red; luminal progenitor-like, slate blue). Numbers within each segment give the percentage of cells analyzed, rounded to the nearest whole percent; the center of each donut is colored by genotype to match the bar graphs below (non-carrier, light brown; BRCA1-carrier, dark brown). BRCA1-carrier PDOs show an expanded luminal progenitor-like fraction (55.6%) compared with non-carrier PDOs (26.2%). Cell types were assigned using the CD49f/EpCAM surface-marker scheme as defined in Methods. Bottom: bar graphs of mean protein expression [log10(Expression + 1)] for CD49f, EpCAM, ER, K14 and K19 in non-carrier (light brown) versus BRCA1-carrier (dark brown) PDOs. EpCAM and ER are significantly elevated in BRCA1-carrier PDOs (**P < 0.01, two-sided Wilcoxon rank-sum test); CD49f, K14 and K19 showed smaller between-group differences. Points represent individual PDOs. Error bars represent standard deviation. **b**, ER expression and cell volume by menopausal status. Both comparisons are restricted to ER-positive cells, that is, cells in which ER was detected above the limit of detection. Donors were classified as pre-menopausal (n = 3), post-menopausal (n = 2) or peri-menopausal (n = 2); the two peri-menopausal donors are not included in this comparison. Left: beehive plots of ER expression [log10(raw AUC)] in pre-menopausal (green) versus post-menopausal (yellow) normal mammary tissue PDOs. Pre-menopausal samples show significantly higher ER expression (****P < 0.0001, two-sided Wilcoxon rank-sum test, n = 498 pre-menopausal cells and n = 236 post-menopausal cells, individual cells as independent observation). Right: beehive plots of cell volume (µm³) for the same ER-positive cells; pre- menopausal cells are significantly larger (****P = 9.89 × 10⁻¹⁶, n = 498 and n = 236 cells). Beehive plots show mean (center line) and standard deviation (whiskers). **c**, PCA of single-cell protein expression. Left: normal mammary tissue PDOs (n = 1,141 cells) colored by cell type (basal-like, coral, n = 620; mature luminal-like, dark red, n = 117; luminal progenitor-like, slate blue, n = 404) with 95% confidence ellipses; PC1 captures 43.8% and PC2 captures 35.7% of variance. Right: normal mammary tissue PDOs colored by menopausal status (pre-menopausal, green, n = 534 cells; post-menopausal, yellow, n = 266 cells) with 95% confidence ellipses, showing relative basal enrichment in post-menopausal PDOs. The right-hand projection includes only the 800 cells from donors of defined pre- or post-menopausal status; the 341 cells from the two peri-menopausal donors shown at left are omitted. Linear arrays of points along PC axes reflect single-protein-positive cells whose PCA projections collapse onto a ray defined by that protein’s loading vector. **d**, PDO individuality scores. Bottom: bar chart of per-PDO individuality scores (diagonal of the k-nearest neighbor similarity matrix, k = 100) for each of the 7 normal mammary tissue PDOs, sorted by decreasing individuality. For each cell, a prior-weighted posterior probability of assignment to each PDO was calculated from its 100 nearest neighbors in three-dimensional CD49f/EpCAM/ER expression space. The individuality score is the mean same-PDO posterior probability across cells from that PDO. Top: schematic illustrating the individuality concept; PDOs with high individuality (e.g., PDO 158)have cells that preferentially neighbor cells from the same PDO, whereas PDOs with low individuality (e.g., PDO 546, PDO 551) have cells whose nearest neighbors frequently originate from different PDO lines.

Among donors with defined menopausal status, ER-positive cells from pre-menopausal PDOs (n = 3 donors; 498 cells) had higher ER AUC values than ER-positive cells from post-menopausal PDOs (n = 2 donors; 236 cells; Fig. 4b). The two peri-menopausal donors were excluded from this comparison. The same ER-positive pre-menopausal cells also had larger measured cell volumes than the post-menopausal cells (mean cell volume 11,391 µm³ versus 8,388 µm³; P = 9.89 × 10⁻¹⁶), and the elevated ER levels were thus at least partly attributable to greater cell volume and total protein content. This distinction, consistent with known hormonal regulation of ER in pre-menopausal breast tissue^25^, highlights cell volume as an important biological variable in single-cell assays where signal scales with total analyte mass^13^. This confound is detectable here only because the CellenONE records a brightfield diameter, circularity and elongation for every dispensed cell, so each AUC measurement is paired with a per-cell volume; assays that load cells stochastically do not retain this pairing. Post-menopausal samples showed relative basal enrichment, consistent with age-related lobular involution that preferentially reduces luminal-rich terminal ductal lobular units^26^.

Principal component analysis (PCA) of single-cell protein expression from normal breast PDOs (n = 1,141 cells) showed partially distinct distributions for the three mammary epithelial lineages, with PC1 and PC2 capturing 43.8% and 35.7% of variance, respectively (Fig. 4c). To quantify inter-donor phenotypic heterogeneity, we computed PDO-level individuality scores based on prior-weighted k-nearest-neighbor voting (k = 100) in 3-dimensional protein expression space (CD49f, EpCAM, ER)^27^ (Fig. 4d). Individuality scores revealed that certain PDOs had greater enrichment of same-PDO cells within their local protein-expression neighborhoods, whereas other PDOs had more extensively overlapping neighborhoods across donors (Fig. 4d). These scores were invariant across a range of k values (k = 80 to 120; SI Fig. 8), indicating that the ranking was insensitive to the selected neighborhood size within this tested range. This analysis demonstrates that AutoBlot’s quantitative, multivariate protein measurements can resolve patient-level phenotypic heterogeneity in cell-limited specimens.

AutoBlot achieves a performance regime unmatched by existing single-cell platforms: approximately 98% targeted-well occupancy and a two-order-of-magnitude reduction in starting-cell requirement from as few as 10,000 starting cells (Fig. 1). This capability is particularly timely as human cell-based models, including organoids, gain regulatory support under the FDA Modernization Act 2.0^28^. We demonstrate that AutoBlot resolves dual ER-immunoreactive bands consistent with ERα66 and ERα46¹³˒¹⁶, which are not discriminated by antibody-binding-only methods such as flow or mass cytometry, quantitatively detects BRCA1-associated differences in targeted protein profiles^15,22–24^, and resolves inter-donor phenotypic heterogeneity across normal breast PDOs using established mammary lineage markers^14,27^ (Figs. 2–4). Because separated proteins are archived in the gel matrix, AutoBlot arrays can be reprobed for additional targets^3^ or excised for downstream LC–MS discovery proteomics. Deterministic co-loading of single cells with single beads further extends AutoBlot to applications requiring precisely paired cell–bead assays, including barcoded single-cell sequencing and proximity-based ligand–receptor binding studies.

## Methods

### PDO culture and dissociation to single cells

PDOs were generated and expanded as described previously^14,20^ under an institutional review board (IRB)-approved protocol. To obtain a single-cell solution, PDOs were digested with TrypLE Express (Gibco, 12604013) for 10 minutes. Digestion was stopped by addition of 10 mL of ice-cold base medium (Advanced DMEM/F12, Gibco, 12634028) with 10 mM HEPES (Gibco, 15630130), 1x GlutaMAX (Gibco, 35050061), 100 U/mL Penicillin-Streptomycin (Gibco, 15140122), and 50 µg/mL Primocin (InvivoGen, ant-pm-05). Cells were centrifuged at 300 x g for 3 minutes and the pellet was resuspended in 100 µL of 1×PBS for subsequent single-cell dispensing into the scWB microwell array.

### MCF-7 cell culture and supplementary scWB analysis

MCF-7 cells were cultured in Dulbecco’s modified Eagle medium (DMEM) supplemented with 5% fetal bovine serum and 1% penicillin–streptomycin. For the experiment shown in SI Fig. 1, scWB arrays were co-immunoprobed with a mouse monoclonal anti-HSP60 antibody (Abcam ab46798) and a rabbit polyclonal anti-GAPDH antibody (Invitrogen, PA1-987), followed by species-specific Alexa Fluor 532-conjugated anti-mouse IgG and Alexa Fluor 488-conjugated anti-rabbit IgG secondary antibodies (Invitrogen), respectively.

### Single-cell western blot assay

The scWB assay was performed using the Milo scWB system (ProteinSimple, Bio-Techne), following the workflow established previously^2,3^. Fabricated in-house, the scWB chips are composed of arrays of 3000 microwells (48 µm diameter × 60 µm height) patterned in a thin photoactive polyacrylamide gel that serves as the assay substrate. The gel functions as both the molecular sieving matrix during polyacrylamide gel electrophoresis (PAGE) and the blotting scaffold during immunoprobing^2^. After cell loading via the CellenONE (see below), chips were inserted into the Milo instrument, which automates the subsequent assay steps: chemical lysis of cells in individual microwells, PAGE for size-based protein separation, and UV-initiated protein immobilization to covalently capture separated proteins in the gel matrix. Lysis was performed for 30 s using a 2× RIPA buffer supplemented with 8 M urea (1% SDS, 0.5% sodium deoxycholate, 0.2% Triton X-100, 2× Tris-glycine), followed by 30 s electrophoresis at 240 V. UV immobilization was performed for 45 s. After immobilization, chips were removed from the Milo, washed, and immunoprobed with fluorophore-conjugated primary antibodies in sequential rounds; round 1: anti-EpCAM-AF647 (0.05 mg mL⁻¹; Biolegend 324212), anti-CD49f-AF488 (0.2 mg mL⁻¹; Biolegend 313608) and anti-ERα-AF555 (0.5 mg mL⁻¹; Abcam ab282199clone SP1, directed against a C-terminal epitope retained in ERα46); round 2 (after stripping): anti-K14-AF488 (0.84 mg mL⁻¹; Novus Bio NBP2-34675AF488) and anti-K19-AF555 (0.5 mg mL⁻¹; Abcam ab203444). All antibodies were incubated for 2 h at room temperature in TBST + 2% BSA, followed by 30 min wash in TBST with buffer exchange every 10 min. Between rounds, antibody probes were stripped for 2 h in a stripping buffer (62.5 mM Tris-HCl (pH 6.8), 2% (wt/vol) SDS, and 0.8% (vol/vol) β-mercaptoethanol). Multiplexed detection relied on sequential rounds of antibody-probe stripping and reprobing^3^. Chips were imaged using a fluorescence microarray scanner (GenePix 4300A; Molecular Devices) with the following settings: CD49f (excitation 488 nm, gain 600, power 50%), ER (excitation 532 nm, gain 700, power 90%), EpCAM (excitation 635 nm, gain 1000, power 100%), K14 (excitation 488 nm, gain 450, power 50%), and K19 (excitation 532 nm, gain 450, power 90%). Protein expression was quantified as described below.

### CellenONE single-cell sorting

Deterministic single-cell loading was performed using the CellenONE robotic dispensing system (Cellenion). The CellenONE uses a piezoelectric dispensing capillary (PDC) coupled with real-time brightfield imaging to identify individual cells in suspension and deposit them one by one into specified microwell positions on the scWB chip^10,11^. An uncoated (Type 0) PDC was used for all experiments. To prevent evaporation of the nanoliter-volume droplets during sorting, the temperature of the base plate was set to track the dew point (+0.5 °C above the measured dew point temperature inside the instrument enclosure). For each scWB slide, 250 cells were sorted, one cell per microwell, into a dehydrated scWB chip (i.e., dried polyacrylamide gel layer), a cell number chosen to demonstrate deterministic single-cell loading; because sorting time is governed by the rate at which cells are identified in the input suspension rather than by the size of the microwell array, additional microwells can be addressed simply by extending the sort duration or raising the input concentration. For routine PDO profiling, 250 cells were dispensed per scWB slide, providing a maximum 95% binomial margin of error of approximately ±6.2 percentage points for estimated cell fractions and a 99.4% probability of sampling at least one cell from a population present at 2% frequency. This required approximately 10 min dispensing time under the conditions used. In the low-input loading demonstration, dispensing 1,000 cells from a suspension at 10,000 cells mL⁻¹ required approximately 45 min; sorting time decreased as input concentration increased toward the upper end of the tested range (30,000 cells mL⁻¹), since cells were identified by the imaging system more frequently per unit time. Cell diameter was recorded by the CellenONE brightfield imaging system during dispensing and used for downstream cell volume calculations. After sorting was complete, the dehydrated scWB chip was placed into a 4-well plate and rehydrated in phosphate-buffered saline (1 min, 1X PBS) prior to loading into the Milo instrument for the scWB assay as described above. For Poisson probability calculation, the nominal microwell volume was calculated from the cylindrical microwell geometry as V=πr^2^h, corresponding to approximately 108.6 pL for a 48-µm-diameter and 60-µm-deep microwell. For stochastic single-cell loading, P(1)=λe^−λ^, where λ=CV. Assuming independent cell and bead loading, the probability of exactly one cell and one bead was calculated as P(1_cell_)P(1_bead_).

### Automated fluorescence image analysis and protein quantification

A custom Python pipeline was developed to automate protein quantification from fluorescence micrographs of scWB chips. The pipeline processes raw fluorescence images acquired after immunoprobing and outputs per-cell protein expression values as area under the curve (AUC) in arbitrary fluorescence units (a.u.). The pipeline proceeds in four stages: In step 1 (image preprocessing), raw fluorescence images were rotated to align microwell rows with the CellenONE deposition sequence, cropped to exclude empty peripheral microwells, and angle-corrected to ensure that electrophoresis lanes were parallel to the image vertical axis. The preprocessed image was then divided into a 10 × 25 grid of regions of interest (ROIs), with each ROI corresponding to a single microwell and abutting electrophoresis lane (SI Fig. 4).

In step 2 (microwell detection and profile extraction), microwell positions within each ROI were identified using the Hough circle transform^29^ with contrast-based validation in which candidate circles were scored by the intensity difference between the dark microwell interior and the surrounding bright gel annulus; candidates failing a minimum contrast threshold of 10 intensity units (8-bit scale) were rejected. A centerline was drawn through the detected microwell center and refined by selecting the x-position (within ±15 pixels of the reference) that maximized the baseline-corrected AUC. The lateral boundaries of a rectangular integration window were defined as ±15% of the ROI width around the refined centerline. A 1D fluorescence intensity profile was then extracted by averaging pixel intensities within this window at each row position along the electrophoresis axis, excluding the microwell region to avoid a dark-pit artifact caused by the microwell void (SI Fig. 5).

In step 3 (baseline correction and AUC calculation), a linear baseline was estimated by performing linear regression on the lowest-intensity fraction (50%) of data points. This baseline was subtracted from the raw fluorescence signal, and the corrected profile was integrated using the trapezoidal rule to yield the AUC, which was carried forward as the quantitative measure of relative protein expression for each cell (SI Fig. 5).

In step 4 (quality control), two quantitative QC gates were applied, followed by array-wide visual inspection. QC 1: the baseline estimation described above ensured robust background fitting. QC 2: a Gaussian function was fitted to the baseline-corrected profile, and lanes with R² < 0.3 were rejected as below the detection threshold. For array-wide visual inspection, a whole-chip overlay of detected microwell positions on the full fluorescence image (green circles for accepted lanes, red circles for rejected lanes) enabled visual confirmation of detection accuracy across the entire scWB array (SI Fig. 7). Manual validation of the ER signal across ten PDO preparations yielded false-positive rates of 0.6–7.4% and false-negative rates of 0.2–14.8% per PDO, with an overall sensitivity of 95% and specificity of 97%. The final output is a tabular file containing lane ID, AUC (a.u.), and cell diameter (µm) for all cells that passed QC, with sub-threshold lanes marked as “below detection limit” (SI Fig. 6).

### Mammary epithelial lineage classification

Cells were assigned mammary epithelial lineage categories independently using surface-marker and cytokeratin measurements. For CD49f/EpCAM classification, cells with detectable CD49f but not EpCAM were classified as basal-like, cells with detectable EpCAM but not CD49f were classified as mature luminal-like, and cells with detectable CD49f and EpCAM were classified as luminal progenitor-like. For K14/K19 classification, cells with detectable K14 but not K19 were classified as basal-like, cells with detectable K19 but not K14 were classified as mature luminal-like, and cells with detectable K14 and K19 were classified as luminal progenitor-like. Cells for which neither marker pair supported a lineage assignment were classified as undefined. Marker detection was determined using the fluorescence-profile quality-control and detection criteria described above.

### Cohort-level protein detection and expression visualization

For each PDO–protein combination, the detection rate was calculated as the number of lineage-classified cells with signal above the detection threshold divided by the total number of lineage-classified cells assayed for that PDO. Mean protein expression was calculated among detected cells after log₁₀(Expression + 1) transformation. For visualization in the bubble matrix, mean expression values were min–max scaled independently for each protein across the seven PDOs. Circle area represents detection rate, whereas circle shading represents the scaled mean expression among detected cells. The 308 cells without an assigned lineage were excluded from this visualization. PDOs were treated as the independent biological units for comparisons associated with germline or menopausal status. Individual cells represent nested measurements within PDOs. Owing to the small numbers of donors, group comparisons are presented descriptively.

### Organoid individuality scoring

To assess whether each PDO possesses a unique, self-contained phenotypic signature or shares protein expression patterns with other PDOs, we adapted the tumor individuality metric from Wagner et al.^27^ using prior-weighted k-nearest neighbor (k-NN) voting in protein expression space. In step 1, the log10-transformed protein expression values for the three markers (CD49f, EpCAM, ER) were standardized to zero mean and unit variance across all cells. A k-NN graph was constructed using Euclidean distance in the standardized 3-dimensional protein expression space, with k = 100 nearest neighbors (excluding self), consistent with Wagner et al.^27^. To incorporate the observed PDO frequencies, in step 2, prior probabilities were computed as W*c* = *n*c / *n*, where *n*c is the number of cells in PDO *c* and *n* is the total number of cells. This prior-weighted formulation followed the tumor-individuality calculation described by Wagner et al^27^. For each cell *i*, the number of its *k* nearest neighbors originating from each PDO *c* was counted, and the posterior probability was calculated as:

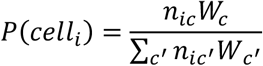

In step 3, a *C* × *C* similarity matrix M (*C* = number of PDOs) was constructed by averaging the posterior probabilities across all cells originating from each PDO: M[*r*, *c*] = mean(*P*(*c*|cell*i*)) for all cells *i* from PDO *r*. The individuality score for each PDO was defined as the diagonal entry of this matrix: Individuality(PDO *r*) = M[*r*, *r*]. High individuality scores indicate that cells from a given PDO predominantly have nearest neighbors from the same donor, whereas low individuality scores indicate that the PDO’s cells share overlapping protein expression profiles with cells from other donors. To assess robustness, the full analysis was repeated for *k* = 80 to 120 in increments of 5; the relative ranking of all PDOs was invariant across the entire tested range, with no rank crossings observed (SI Fig. 8).

### Use of AI-assisted tools

Claude (Anthropic) was used during manuscript preparation for two purposes: (1) assisting with text clarification and drafting revisions in response to co-author feedback, and (2) debugging Python scripts for data analysis. All scientific content, data interpretation, experimental design, and conclusions were produced by the authors. The AI tool was not used for data generation or independent analytical decision-making.

## Software and code availability

All image analysis and computational scoring pipelines were implemented in Python using NumPy, pandas, scikit-learn (StandardScaler, NearestNeighbors), and matplotlib. Custom scripts for fluorescence image processing, baseline correction, Gaussian QC filtering, and PDO individuality scoring are available at https://github.com/herrlabucb/autoblot-scwb-auc.

## Author contributions

Conceptualization: CD, NI, NG, JMR, AEH

Formal analysis: CD, NI

Funding acquisition: AEH, JMR

Investigation: CD, NI, NG, LJ, JMR, AEH

Methodology: CD, NI, NG, LJ, JMR, AEH

Software: CD, NI

Supervision: AEH, JMR

Visualization: CD, NI

Writing – original draft: CD, NI

Writing – review & editing: CD, NI, NG, JMR, AEH

## Acknowledgments

We thank Joshua Cantlon (Scienion) for helpful discussions and technical support related to the CellenONE platform.

