## Supplemental Files for "AutoBlot: deterministic single-cell western blotting reveals proteomic diversity in rare cell populations"

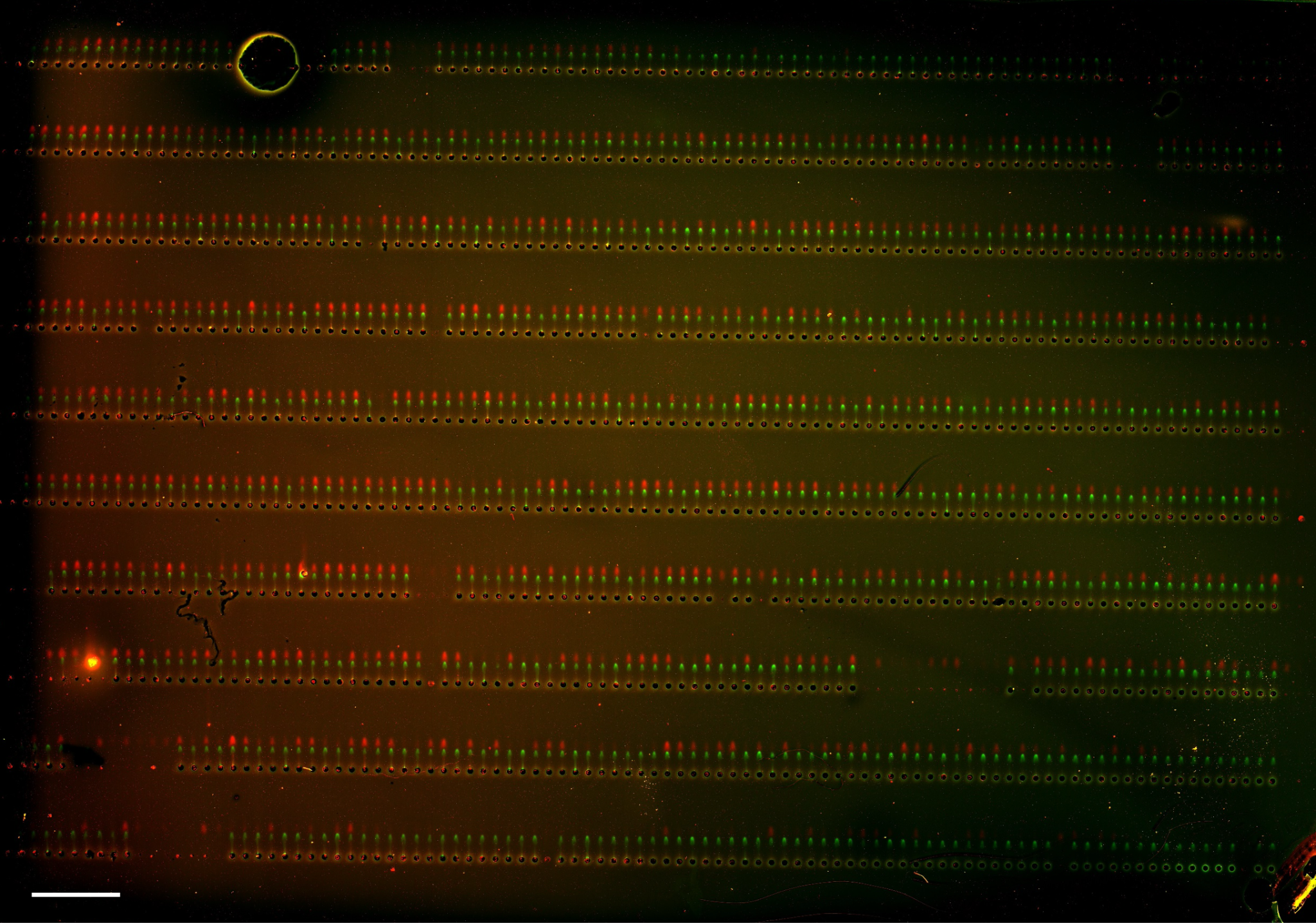

SI Fig. 1: Representative false-color fluorescence micrograph of an AutoBlot microwell array following single-cell Western blotting of MCF-7 cells. Targeted microwells (48 µm diameter, 60 µm height) contain individually dispensed cells that was lysed in situ, with proteins separated by electrophoresis and immobilized via UV-activated photocapture. The array was probed for HSP60 (red) and GAPDH (green). Distinct fluorescent bands are visible at expected electrophoretic migration distances from the microwell, confirming successful protein separation and detection at the single-cell level. Starting from an input of only 10,000 cells, AutoBlot captured approximately 1,000 individual cells across the array, corresponding to approximately 10% recovery of the starting cell input. Among microwells targeted for deposition, approximately 98% contained a single cell. Both HSP60 (mitochondrial chaperonin, ~60 kDa) and GAPDH (glycolytic enzyme, ~36 kDa) are robustly detected across the majority of occupied microwells, demonstrating consistent lysis, electrophoretic separation, and immunoprobing performance at scale. The high density of positive signals across the array underscores AutoBlot's ability to generate statistically powered single-cell proteomic datasets from limited sample inputs, a critical requirement for clinical specimens such as patient-derived organoids or fine-needle aspirates where cell numbers are inherently constrained. Scalebar: 1 mm.

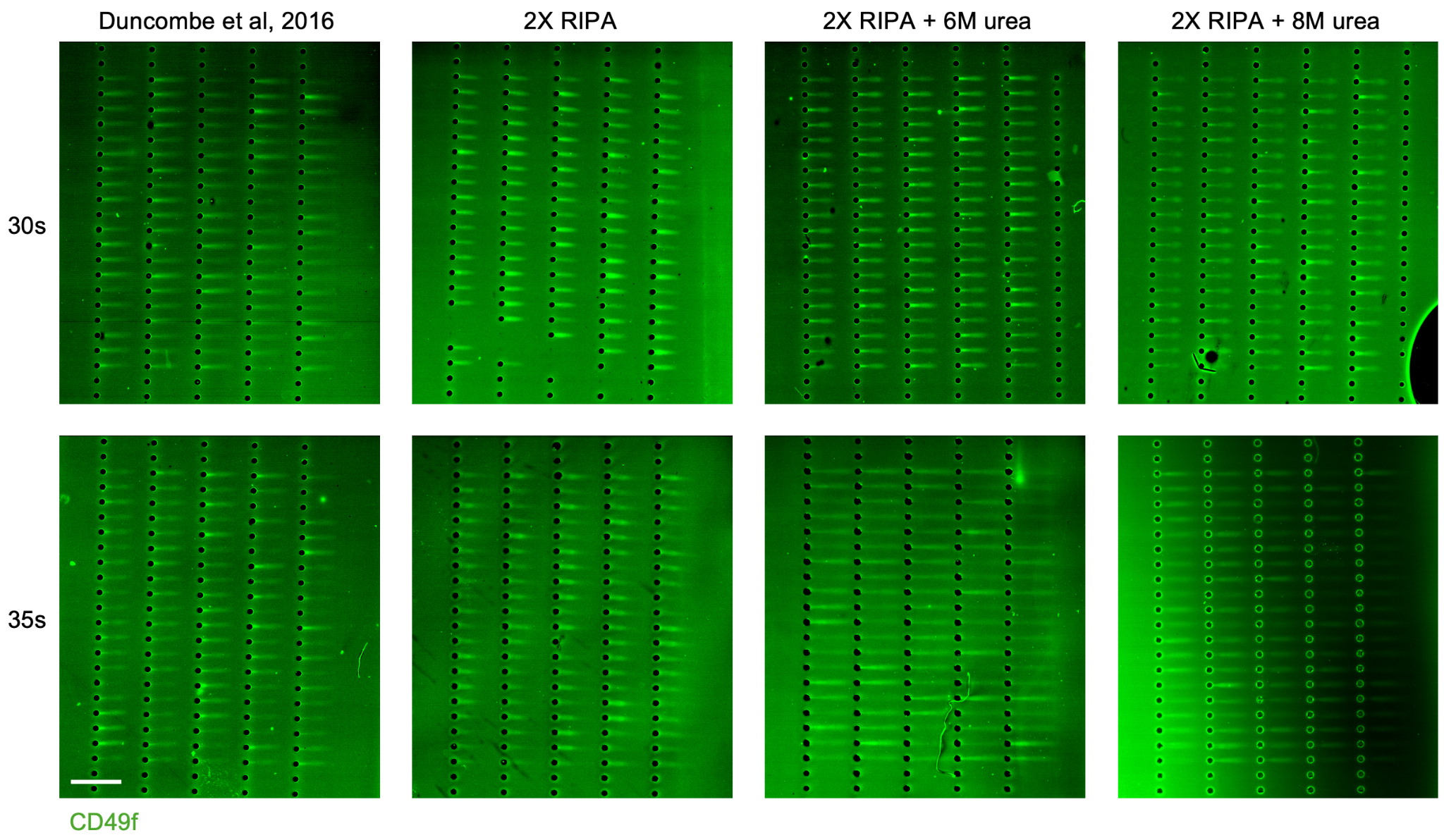

SI Fig. 2: Optimization of lysis buffer composition and electrophoresis duration for single-cell Western blotting of CD49f in patient-derived breast cancer organoid, comparing the standard Duncombe et al. 2016 formulation against 2X RIPA, 2X RIPA + 6M urea, and 2X RIPA + 8M urea at 30 s and 35 s electrophoresis times. The PDOs used in SI Figs. 2 and 3 were independent optimization samples and were not included in the seven-donor normal-tissue cohort. Scalebar: 1 mm.

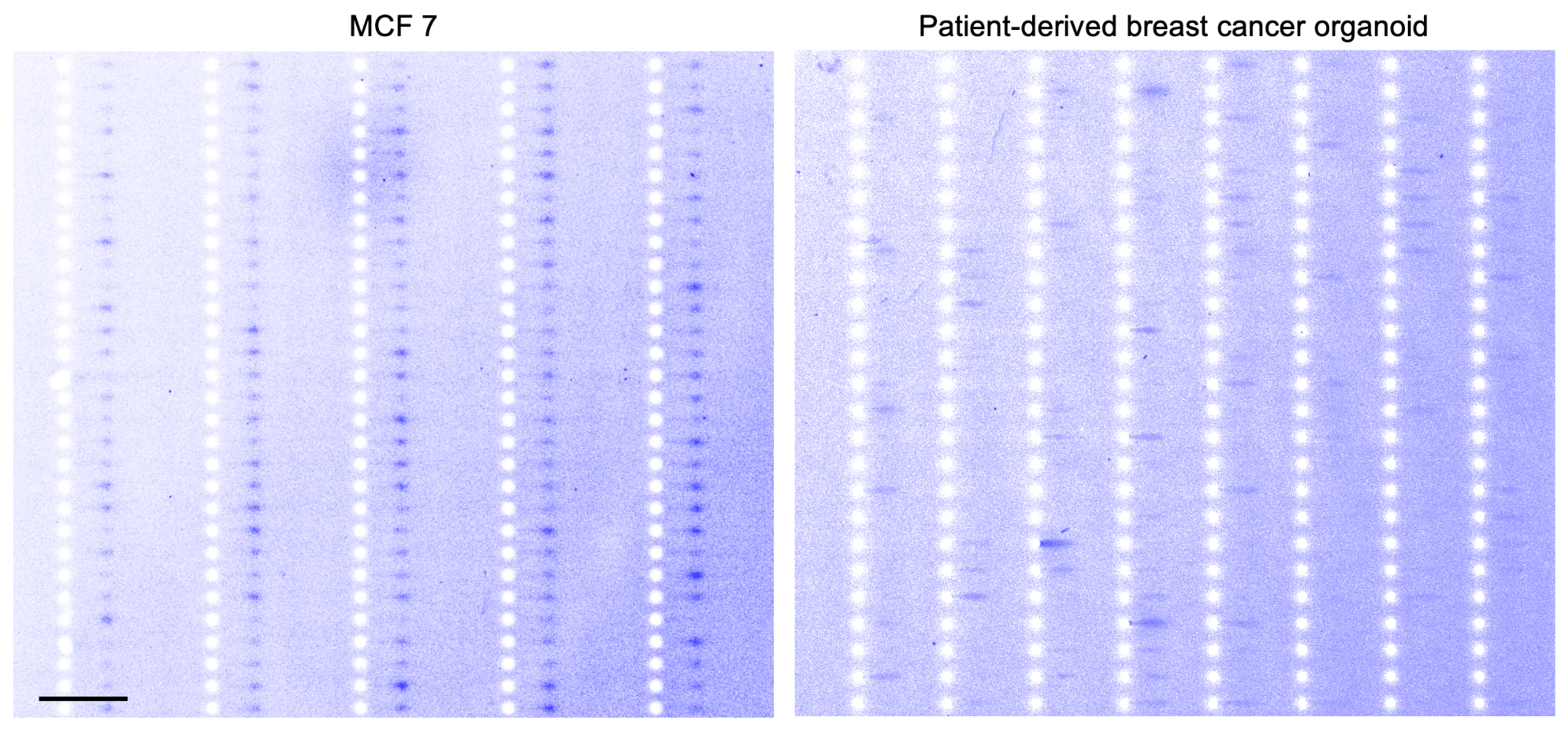

SI Fig. 3: Patient-derived breast cancer organoids preserve single-cell protein heterogeneity in EpCAM expression that is lost in the established MCF-7 cell line, as resolved by AutoBlot single-cell Western blotting. Scalebar: 1 mm.

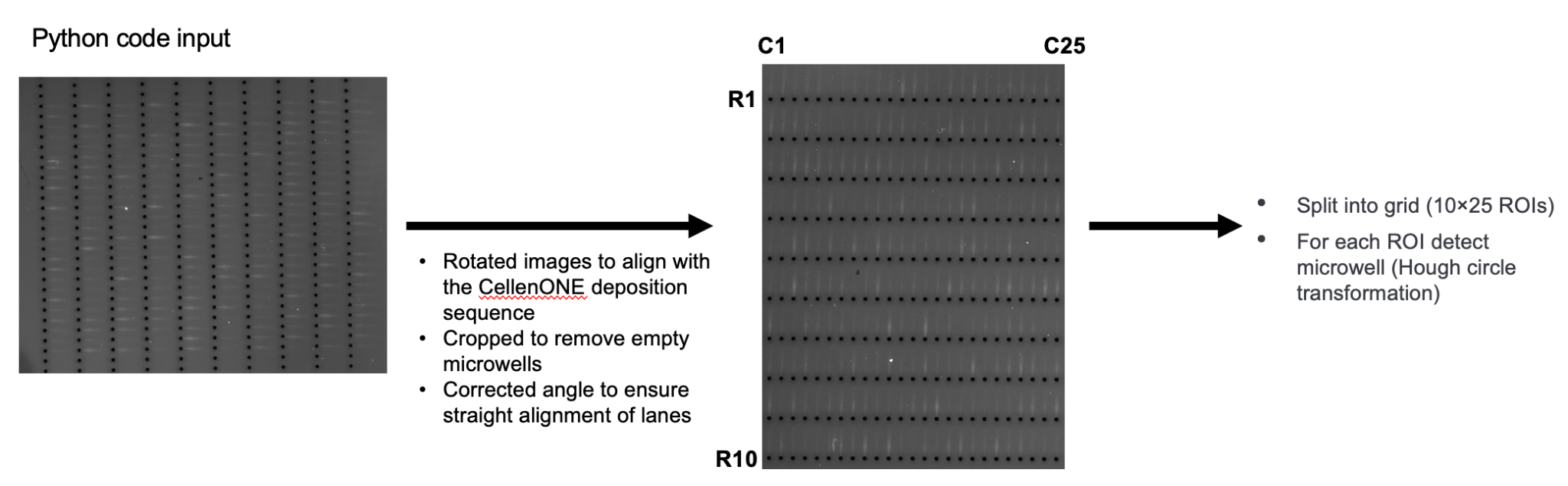

SI Fig. 4: Automated image processing pipeline for single-cell Western blot quantification: raw fluorescence micrographs are rotated to align with the CellenONE deposition sequence, cropped, and segmented into a 10×25 grid of individual lane ROIs, with microwell positions detected within each ROI using Hough circle transformation and contrast-based validation.

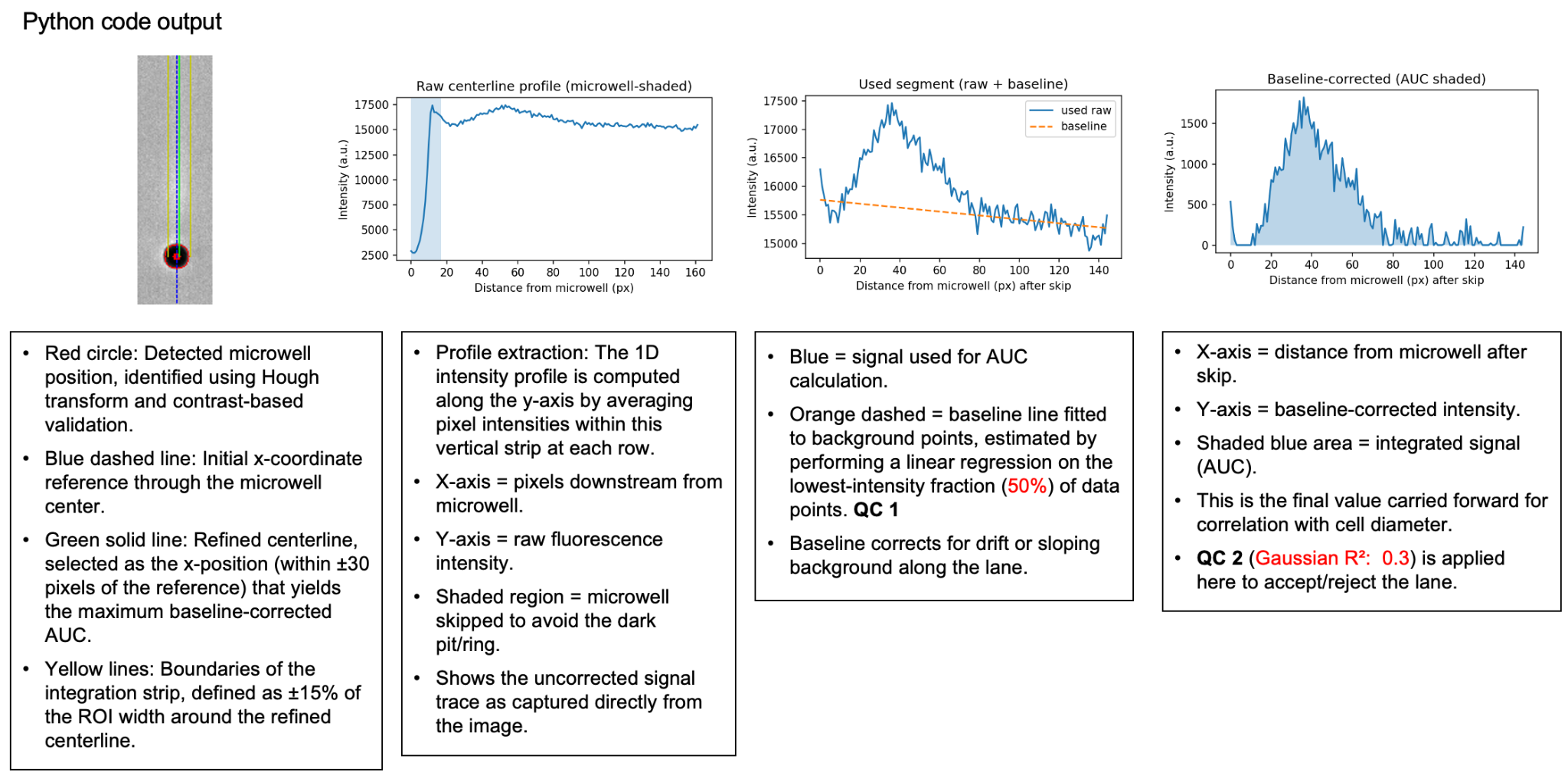

SI Fig. 5: Per-lane signal extraction and quantification workflow: a 1D intensity profile is sampled along a refined centerline through each microwell, the microwell region is excluded, a linear baseline is fitted to the lowest 50th percentile of intensities (QC 1), and the baseline-corrected profile is integrated to yield the area under the curve (AUC), with a Gaussian R² ≥ 0.3 threshold applied as a second quality control gate (QC 2) to accept or reject each lane.

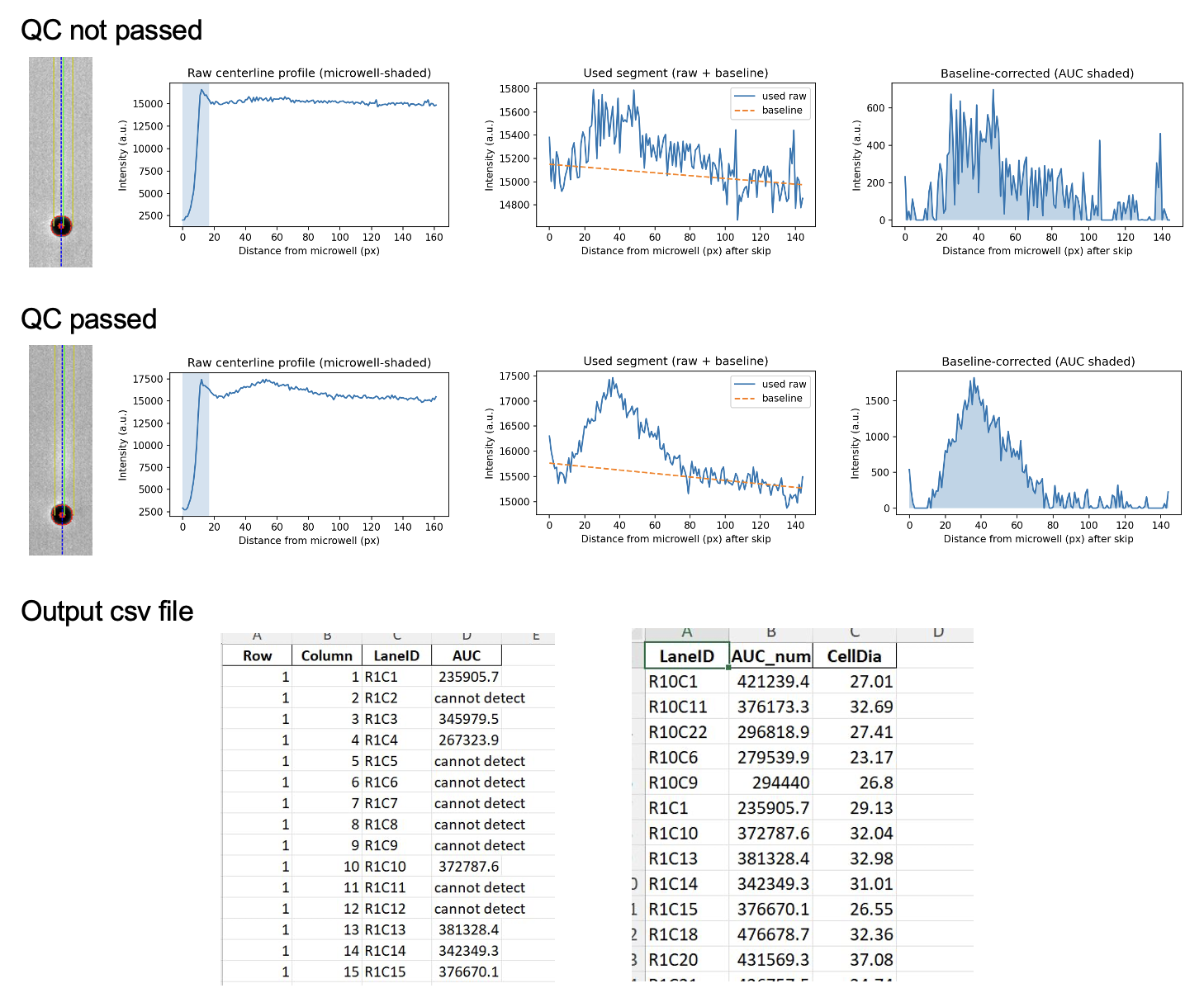

SI Fig. 6: Representative examples of lanes that failed (top) and passed (bottom) the two-stage quality control, alongside the output csv files reporting AUC values in arbitrary units for all detected lanes and the matched AUC with cell diameter for lanes passing QC.

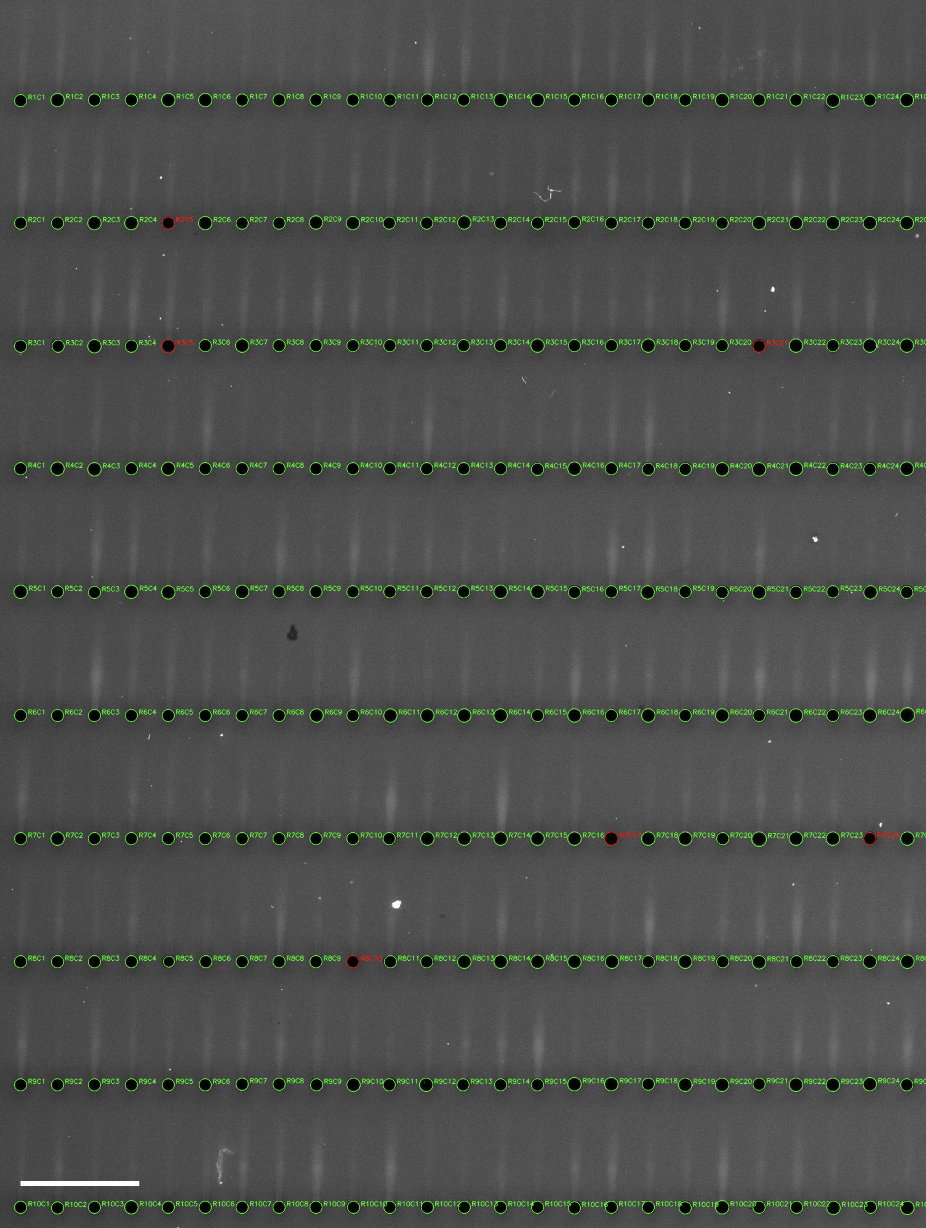

SI Fig. 7: Full-array quality control overlay showing all detected microwells annotated with lane identifiers, where green circles indicate lanes that passed both QC gates and red circles indicate lanes that failed, enabling rapid visual inspection of array-wide signal quality. Scalebar: 1 mm.

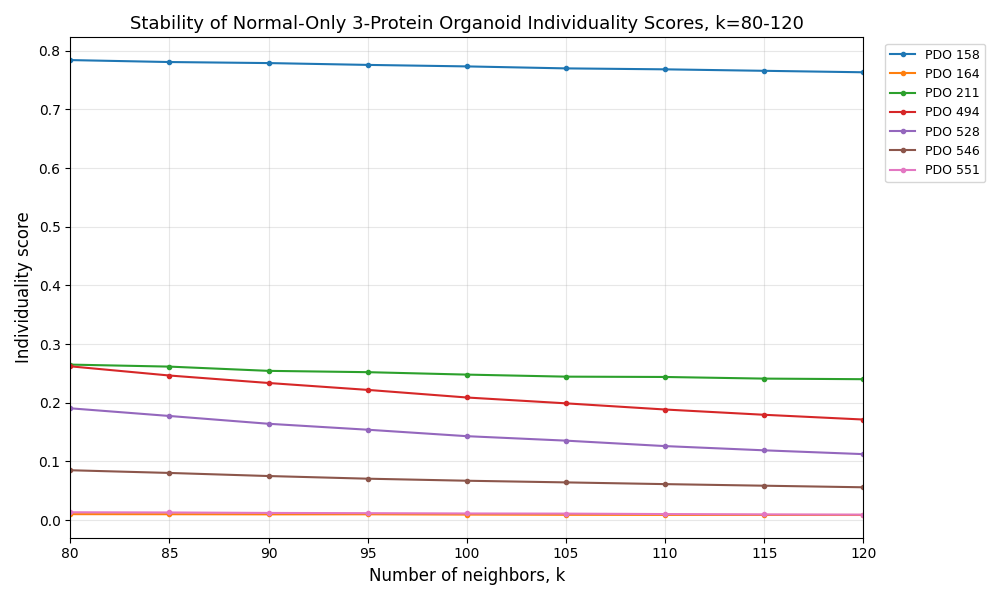

SI Fig. 8: Organoid individuality scores, computed via k-nearest neighbor analysis of single-cell protein expression profiles, remain rank-stable across a range of k values (80 to 120), confirming that inter-organoid proteomic differences are robust to parameter choice.

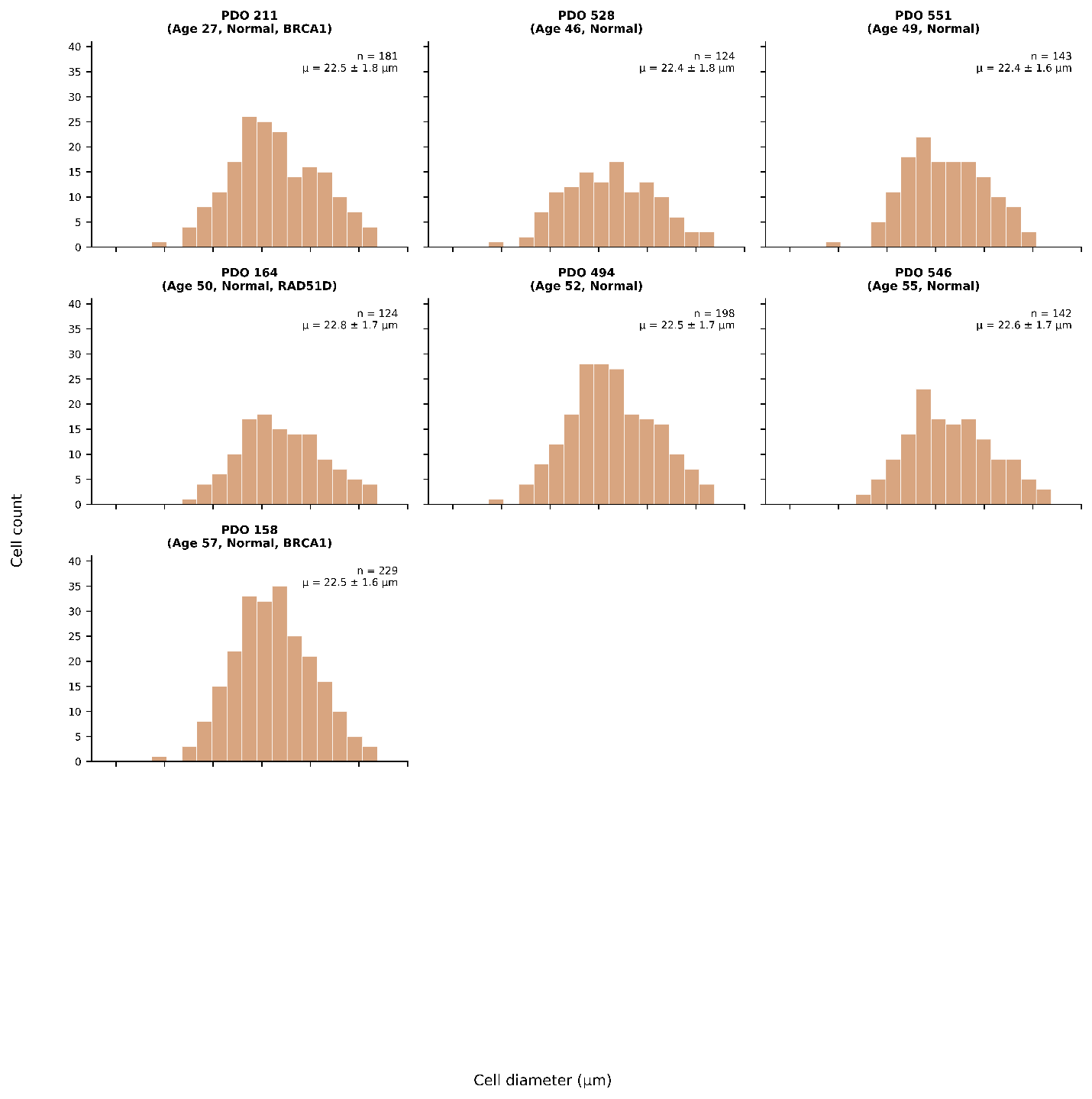
Fig. 9: Cell diameter distributions measured by CellenONE brightfield imaging during single-cell dispensing are consistent across 7 patient-derived organoids, with mean diameters ranging from 21 to 24 µm.
